# A Conserved Mechanism in Eye Optical Development: Lens Nucleus Centralization in *Xenopus laevis*

**DOI:** 10.1101/2025.11.09.687506

**Authors:** Karla A. Garcia, Kelly Ai-Sun Tseng, Irene Vorontsova

**Author notes:** Co-corresponding author Contact: Kelly Tseng. Co-corresponding author Contact: Irene Vorontsova.

## Abstract

Developing eye optics, determined by the lens and cornea, must coordinate with the axial length of growing eyes to focus light onto the retina to form an image. It was found that zebrafish (*Danio rerio)* lens nuclei are initially anteriorly localized in optical axes in larvae, then centralize at older stages. An anteriorly placed lens nucleus would increase lens power, thereby likely enabling a functional optical system in larvae, where eye axial length is short. To assess if alike mechanisms occur in other aquatic animals, we studied the clawed frog, *Xenopus laevis*, a fully aquatic species similarly relying on vision for survival at stages where eyes are small. We found the *Xenopus* tadpole lens nucleus also shifted from an anterior to a central location in the optical axis during the prometamorphosis period. Similarly, in eyes regenerated after embryonic ablation, tadpole lens nuclei are anteriorly localized then centralize before metamorphosis, recapitulating the same pattern as control developing eyes. Moreover, lens nuclei localization in optical axes in developing and regenerated *Xenopus* eyes show close correlations to axial eye length. Close correlation of these two parameters suggests lens nuclei centralization is required for a functional optical system by coordinating the focal length. Our findings suggest a conserved evolutionary mechanism for eye optical development in at least two aquatic species. Understanding key mechanisms regulating crosstalk between eye optics and eye axial length will aid in discovering mechanisms of optical development and future therapies to prevent or delay formation of refractive error when these two properties mismatch.

## Introduction

The transparent lens of the eye transmits and focuses light on the neural retina and is crucial for providing high resolution vision. In aquatic species, the ocular lens is the main optical element for light refraction, and its shape is generally observed to be spherical (Sivak, 1988), which together with a higher maximum refractive index compared to terrestrial lenses (Kroger, 2013), enables higher optical power. The refractive index, protein expression and physiology change dramatically as a function of lens depth, which also represents the age gradient. The central lens fiber cells make up the lens nucleus, which has the highest refractive index, and its relative location and optical properties are essential for overall lens power and optical quality. Although generally described to be centrally localized in the optical axis, the lens nucleus can change its location during optic development in at least one species.

The nucleus of an excised larval zebrafish (*Danio rerio*) lens is anterior localized in the optical axis, which centralizes to a central location in the optical axis of the lens at mid juvenile stages, ∼ 15 mm standard length (Vorontsova et al., 2018). Since zebrafish require functional vision from around 5 days post fertilization (dpf) when their yolk depletes, it has been shown that this coincides with a well-developed visual system (Fleisch and Neuhauss, 2006). We proposed that an anteriorly positioned lens nucleus allows zebrafish larvae to focus light onto the closely adjacent retina at these young stages. As the eye increases in size during development, the lens nucleus centralizes, thereby increasing optical power to focus light onto the retina of the growing eye. The centralization of the lens nucleus must also be matched with the developing refractive properties, which also reach a maximum value at same stages of development ∼ 15 mm standard length (Wang et al., 2020). The mechanism of lens nucleus centralization has not been identified in other species of mammalian or aquatic species before; thus it is not known whether this mechanism is shared with other aquatic animals or is unique to this species.

The African clawed frog (*Xenopus laevis*) is a fully aquatic species that is a widely used biomedical research model and has well-characterized eye and lens development processes that are similar to mammals (Viet et al., 2020). *Xenopus* embryos have rapid external development, are largely transparent, and thus highly accessible for studies of ocular development. Notably, *Xenopus* embryos exhibit robust eye regrowth capabilities during tailbud embryos stages that is age-dependent (Kha et al., 2019; Kha et al., 2018). It is also capable of regenerating individual ocular structures including the retina, optic nerve, and lens (Gaze, 1959; Ide et al., 1984; Rio-Tsonis and Tsonis, 2003; Tseng, 2017). In contrast, zebrafish readily regenerates its retina (Wan and Goldman, 2016), RPE (Leach et al., 2021), optic nerve (Pérez-Montes et al., 2026) but appear to lack lens regeneration ability (Suetsugu-Maki et al., 2012; Tsonis et al., 1993). Because of its suitability and accessibility, *Xenopus* is ideal for studies of optical physiology but has been underutilized.

It is possible that *Xenopus* also use centralization of the lens nucleus to modulate optical power during development. Here, we show that developing *Xenopus laevis* lenses, like *Danio rerio*, undergo centralization of the lens nucleus from an initially anterior localization in the optical axis. Furthermore, the lens nuclei of regenerated eyes following optic cup removal at the mid-tailbud stage (Nieuwkoop-Faber st. 27) recapitulates centralization pattern of developing eyes through to adulthood. The normalized localization of the lens nucleus strongly correlates with eye axial length, indicating tight coordination between these two parameters. This study suggests lens nucleus centralization is a conserved mechanism required for vision of these two aquatic animals. Our work also provides a new platform for comparative studies of visual function and optical physiology in developing versus regenerating eyes.

## Methods

The animal protocols used in this study adhere to and have been approved by the Institutional Animal Care and Use Committee (IACUC) of the University of Nevada, Las Vegas and adhered to the ARVO Statement for the Use of Animals in Ophthalmic and Vision Research. Embryos were obtained using *in vitro* fertilization and were cultured in 0.1X Marc’s Modified Ringer (MMR, 0.1 M NaCl, 2.0 mM KCl, 1 mM MgSO4, 2 mM CaCl2, 5 mM HEPES, pH 7.8) medium. Staging was done according to the Normal Table of Xenopus Laevis (Daudin)(Faber, 1994). Embryonic eye ablation surgery was performed as previously described (Kha et al., 2018). Briefly, st. 27 embryos were anesthetized using tricaine. The left eye was carefully removed using Dumont No. 5 forceps with the uninjured contralateral eye served as the control. After surgery, embryos were washed, allowed to recover in 0.1x MMR and cultured at 22º C. Fully regrown eyes containing normal size and morphology (Kha et al., 2018) were analyzed in this study.

For eye and lens assessment, euthanasia was performed with either 0.5g/L tricaine (Tokyo Chemical Industry, Japan) for tadpoles or 4% benzocaine for adult frogs. The eyes and lenses were excised and imaged in 1X Phosphate Buffered Saline using a protocol previously described for zebrafish (Vorontsova et al., 2018; Vorontsova et al., 2019). Lens and eye images were obtained using a Zeiss V20 stereomicroscope with an AxioCam MRc camera (Jena, Germany). Whole eyes were imaged in the axial orientation (perpendicular to the optical axis) and measurements taken from the cornea to the back of the eye approximately through the optical axis. Lens nucleus localization was measured in both, axial (perpendicular to the optical axis) and equatorial (through the optical axis) orientations as described previously (Vorontsova et al., 2018; Vorontsova et al., 2019). Measurements were made using ImageJ software (version 1.54k, 15 September 2024) to express the nucleus location as the normalized localization of the lens nucleus with respect to the lens radius, r/a, where *r* is the distance from the center of the lens to the center of the nucleus and *a* is the lens radius. An r/a value of 0.0 indicates central localization of the lens nucleus in the lens along the optical axis, while values larger than 0.0, are closer to the anterior pole in lenses in axial orientation. Values of r/a closer to 1.0 indicate the lens nucleus is closer to the pole. Values of r/a over 0.0 in equatorial planes indicate the localization of the center of the lens nucleus to the closest side.

## Results

*Xenopus laevis* lens development is highly similar to mammals (Viet et al., 2020). At the early swimming tadpole stage, st. 44, the lens nucleus is formed and is composed of primary fiber cells. The adult lens structures are present by st. 48. The eye grows proportionally in size (Beach and Jacobson, 1979; Wen and Shi, 2016) as the tadpole continues to grow through pre-metamorphosis (st. 45-54), pro-metamorphosis (st. 55-58), and metamorphic climax (st. 58-66). The clawed frog, like zebrafish, have functional vision at young stages of development, when the eye is small and lens to retina distance is minimal. *Xenopus* is a fully aquatic anuran post metamorphosis, therefore has a spherical lens like fish (Mitra et al., 2022).

We examined the *Xenopus* lens nucleus localization pattern starting from st. 44 through to adulthood, during which time the lens diameter increased from ∼140 µm to ∼1200 µm. At young tadpole stages, lenses have lens nuclei localized closer to the anterior pole (Fig. 1) with the highest mean r/a value is approximately 0.45, and maximum 0.5 (Figure 1A, triangles). This asymmetrically localized lens nucleus is evident in representative lenses from a stage 50 tadpole shown in the axial orientation (Figure 1B). The location of the lens nucleus gradually centralizes during development to reach a plateau at a mean normalized r/a ∼0.08 in the optical axis at about st. 60 (Figure 1A triangles). This indicates that the lens nucleus stops centralizing just short of the very center of the optical axis of the lens. A representative lens is shown for st. 64 with the lens nucleus appearing centrally localized in the optical axis (Figure 1D). The same lenses were also analyzed for lens nucleus location in the equatorial orientation, which is directly through the optical axis. The center of the lens nucleus is central in this orientation at all developmental stages examined in this study (Figure 1A, circles) with representative lenses oriented equatorially at young (Figure 1C) and old lenses (Figure 1E) displaying centrally localized lens nuclei.

**Figure 1.**
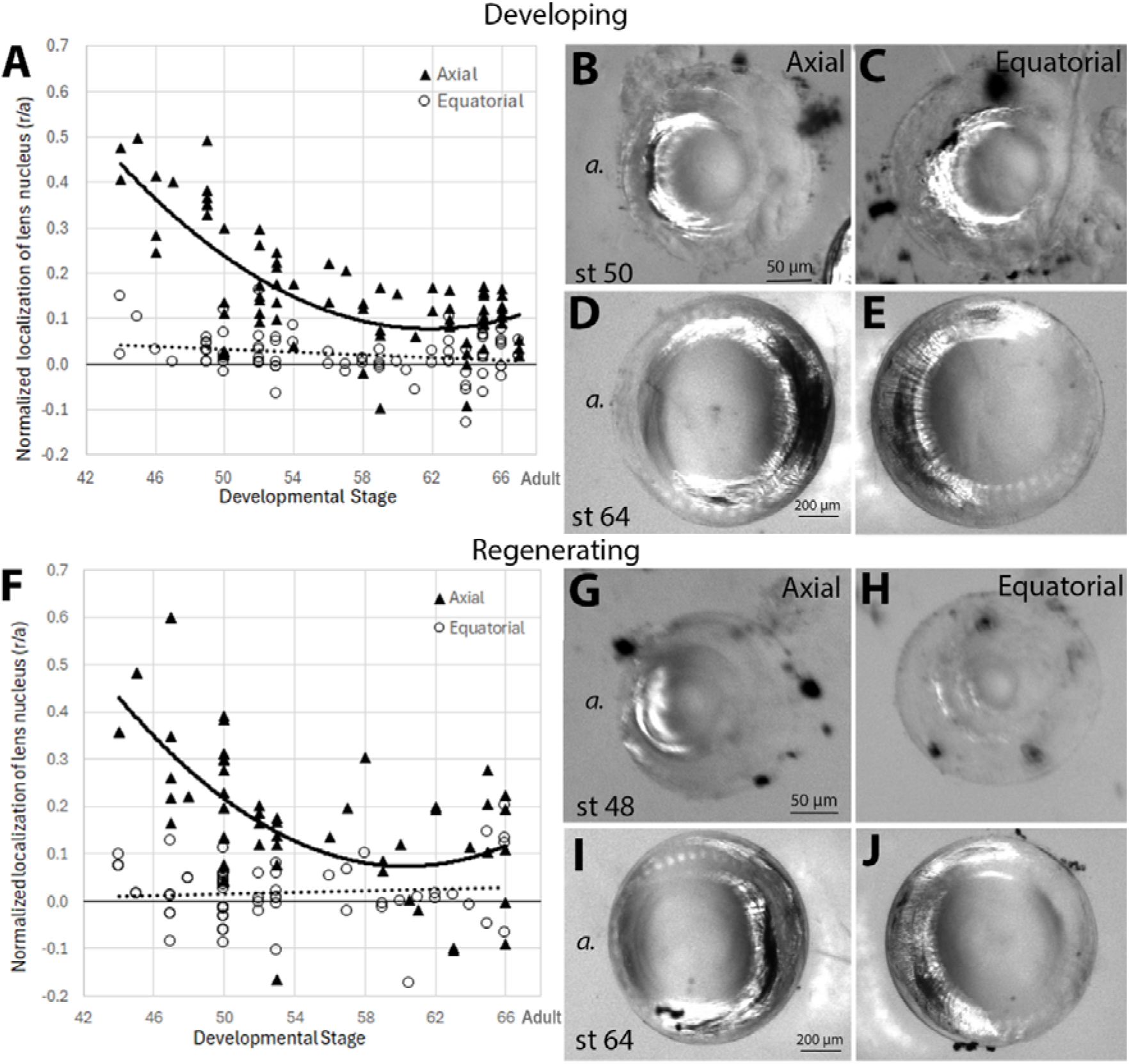
Xenopus laevis lens nucleus centralizes in the optical axis from an initial anterior location. Lenses from developing (**A-E**) or regenerated eyes (**F-J**) were imaged either perpendicular to the optical axis (axial orientation, **B, D, G, I** a.=anterior to the left) or through the optical axis (equatorial orientation, **C, E, H, J**). Normalized lens nucleus localization measured in an axial orientation (developing **A**, polynomial fits R^2^=0.5481, n= 74 and regenerating **F** R^2^=0.4092, n=61), and equatorial orientation (developing **A**, linear fit R^2^=0.0529, n=61; and regenerating **F** R^2^=0.0081, n=59). The distance of the center of lens nucleus (r) centralizes from an anterior localization (higher value of r/a) as a function of the of the lens radius (a) in developing (**A, triangles**) or regenerating lenses (**F, triangles**) but is always centrally localized in equatorial orientation (**A, F circles**).

At st. 27, the embryonic *Xenopus* eye contains the lens placode and an optic cup. After ablation of the embryonic eye, the embryo can fully regenerate a functional eye within 5 days (∼st. 44-45) that is morphologically indistinguishable to an uninjured eye and recapitulates the overall retinal differentiation process (Kha et al., 2019; Kha et al., 2018). Both cellular and molecular mechanisms have been characterized (Guerin et al., 2025; Hack et al., 2024; Kha et al., 2019; Kha et al., 2023). We took advantage of this unique *Xenopus* capability to examine the pattern of lens nucleus localization in fully regenerated eyes. A very similar pattern to developing lenses is observed in fully regenerating lenses at larval stages (Figs. 1F-J). In young tadpoles, a normalized lens nucleus localization mean r/a value of ∼0.45 at st. 44 is observed and a maximum value of 0.6 observed at stage 47. The predicted trend value or r/a plateaus at ∼0.08 at st. 59-60 (Figure 1F, triangles). Representative lenses show that the lens nucleus is anteriorly localized as shown for st. 48 (Figure 1G) and it is central as development progresses, as shown for st. 64 (Figure 1I). In the equatorial orientation, localization of the lens nucleus is averages an r/a of close to 0.0 at all stages (Figure 1F circles), and is shown in the representative lenses (Fig. H and J).

A direct comparison of lens nucleus localization patterns in developing and regenerating *Xenopus* eyes revealed a strong correlation in lens nucleus centralization mechanisms between developing and regenerated lenses (Figure 2A), which look almost identical throughout development (from st. 44 to froglet). To assess whether lens nucleus localization also correlates with eye axial length in regenerated eyes, we first examined the relationship between lens diameter and axial eye length in regenerating eyes as any differences between these parameters would affect the optical system. In both developing and regenerating eyes, tight linear correlations between the axial diameter and axial eye length of the lens were observed (Fig. 2B, r^2^=0.8544 for developing as compared to r^2^=0.9383 for regenerating eyes). This indicates that in fully regenerated eyes, axial eye length matches that of normally developing eyes by st. 44, and lens size correlates to the axial eye length also. We further hypothesized that the anteriorly placed lens nucleus is required to match the axial eye length of eyes at younger stages to facilitate a functional optical system. To test this, we compared axial eye lengths as imaged perpendicular to the optical axis with the lens nucleus localization r/a values from the same eye in axial orientation (Figs. 2C-F). This comparison shows a very similar relationship between r/a of lens nucleus localization and the axial eye length in developing and regenerated eyes (Figure 2G). For both developmental pathways, the lens nucleus is closer to the anterior pole in smaller eyes and is more central in larger eyes.

**Figure 2.**
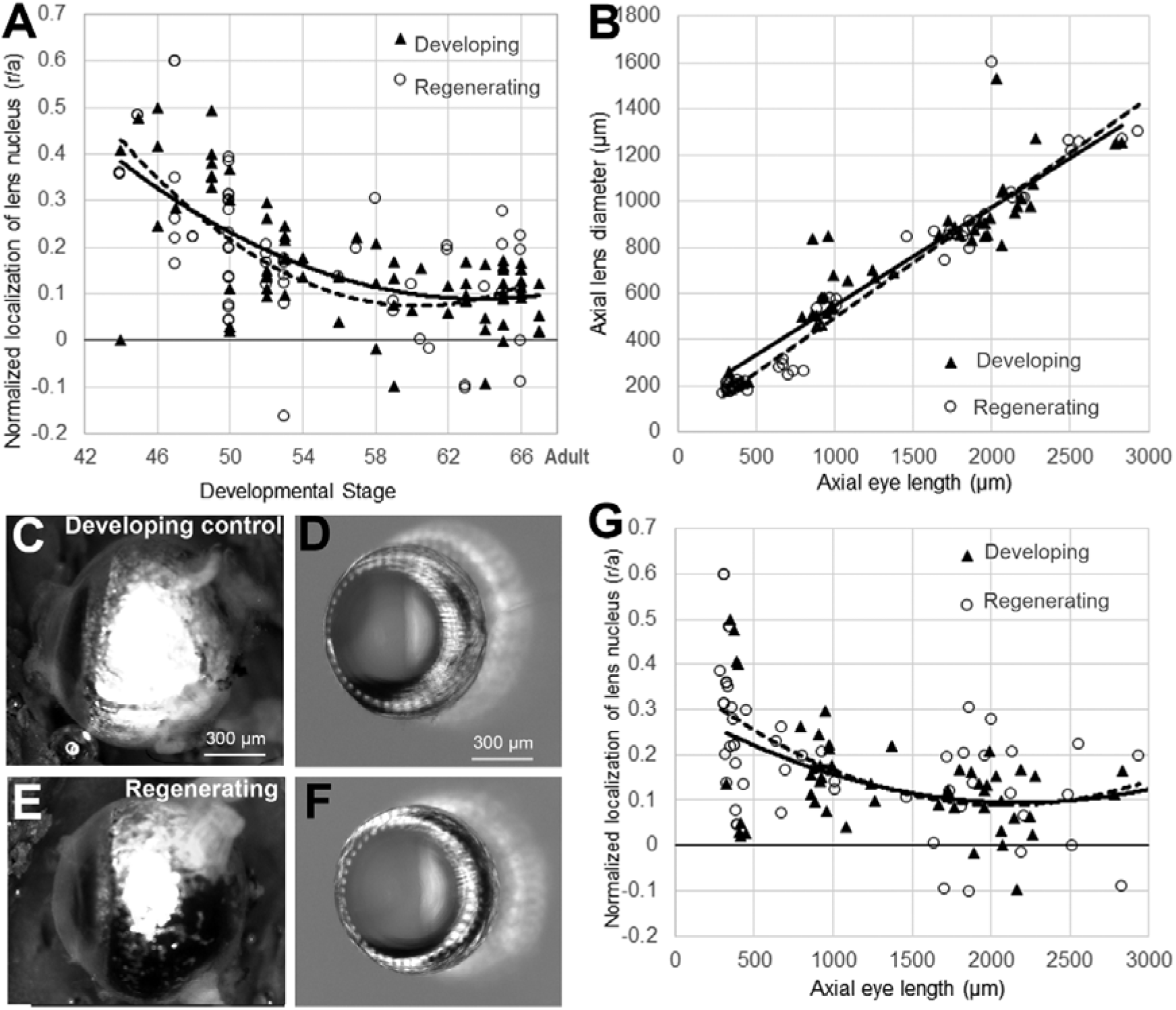
Lens nucleus centralization correlates with axial eye length in developing and regenerating lenses. Lens nuclei follow a similar trend of centralization from stage 44 in developing (**A triangles**, polynomial trend r^2^=0.4092, n=68) and regenerating lenses (**A circles**, polynomial trend r^2^=0.4804, n=-61) reaching a plateau of r/a of ∼0.1 at stage 58. Strong linear correlation between axial lens diameter and axial eye length was observed (**B** n= 77, r^2^=0.8544 for developing (**triangles**) and n=63, r^2^=0.9383 for regenerating (**circles**) eyes). Example of a developing eye at stage 60 (**C**) and it s excised lens (**D**) with anterior to the left. Example of a regenerating eye (**E**) at stage 60 with its excised lens (**F**) with anterior to the left. The centralization of the lens nucleus closely correlated to axial eye length in both, developing (**G triangles**, polynomial trend r^2^=0.2026) and regenerating (**G circles**, polynomial trend r^2^=0.3251) eyes.

## Discussion

The developmental centralization of the *Xenopus laevis* lens nucleus in the optical axis from an initial anterior position, as previously observed in *Danio rerio*, supports the requirement of this mechanism for functional optical development to increase visual acuity at young stages. The conservation of this mechanism between at least two fully aquatic species suggests that it is likely present in other aquatic species with similar requirements for vision. For light to focus on the closely located retina of the larval zebrafish or *X. laevis*, an anteriorly placed lens nucleus would facilitate a higher power lens to enable light to focus onto the photoreceptors. During *X. laevis* development from st. 44 to adult, the eye axial length increases more than 6-fold, which would require a dynamic change in focal length to enable an emmetropic eye. Furthermore, our results indicate that embryonically regenerated lenses of *X. laevis* undergo the same process of lens nucleus centralization that is closely linked to axial eye length, which recapitulates the normal developmental program. The close correlation between lens nucleus location and axial eye length in both developing and regenerated eyes provides evidence that these two parameters are tightly coordinated to produce optical power of the lens to match the axial length of the eye to focus light onto the retina of the growing eye. This raises the idea that the development of optical properties of the anterior eye, mainly the lens, are modulated by the growing eye.

Although mammalian lenses to date have not been specifically examined for similar lens nucleus asymmetry at early stages of development, there have also been no reports of this phenomenon. Other studies assessing aspects of lens anatomy during mouse prenatal and postnatal stages that do not display asymmetrically placed lens nuclei in the optical axis (Aoki et al., 2016; Cheng et al., 2016). The apparent absence of this mechanism could be due to a difference in requirements for functional vision at infancy to avoid predators and secure food. *X. laevis* (Liu et al., 2016) and zebrafish (Gestri et al., 2012), like many other aquatic animals (Pankhurst and Eagar, 1996) greatly rely on vision for sensory input at young/larval stages. Land mammals vary in the immediacy of their reliance on vision with mammals including humans, primates, most rodents and others that open eyes, and mature their eye anatomy and visual processing pathways well after birth and into childhood (Danka Mohammed and Khalil, 2020). Precocial animals, however, have functional vision when they are born or hatch, including ground nesting birds such as chickens (Álvarez Hernán et al., 2021), ducks, and ungulates (grazers) (Hamor and Whitley, 2024). This contrasts with a requirement for functional vision due to the independence needed for feeding and evading predators for larval zebrafish and *Xenopus*. Interestingly though, it has been found that in normal aging human lenses, there are asymmetric changes in optical properties that results in the age-related hyperopic shift in lens power (Lie et al., 2018). This does suggest that shift of the high-refractive index lens in the optical axis may be a mechanism for regulating lens optical power in animals including humans.

Besides behavioral differences, anatomical constraints in aquatic eyes, including short vitreous chambers of aquatic animals at younger stages and almost spherical lenses all stages of development in fish and *X. laevis* could drive the centralization of the lens nucleus to achieve functional vision at different stages. We propose that lens nucleus centralization serves two purposes: 1) Increase lens power at young stages for a functional optical system to focus light onto the closely adjacent retina, and 2) As a tool for developing lens optical power that is analogous to dramatic lens shape change in human lenses during childhood (Tripathi and Tripathi, 1983).

In land vertebrates, the cornea provides about two thirds of optical power, while the lens accounts for the remaining third (Land and Fernald, 1992), however it is the lens that modulates its optical power to achieve an emmetropic eye (Mutti et al., 2018). In fully aquatic species, such as zebrafish or *X. laevis*, the high refractive spherical lens contributes majority of optical power (Greiling and Clark, 2008; Land and Fernald, 1992) with minimal contribution from the cornea. There are two phases in emmetropization in humans. The first being exponential growth up to 2 years of age, followed by a slower phase of eye growth that is primarily compensated by changes in lens optical power (Mutti et al., 2018) in which visual input guides eye growth (Summers et al., 2021). The regulation of eye growth by visual input has also been reported in many other species, including fish (reviewed in (Troilo et al., 2019)). This mechanism has been instrumental in developing clinical treatments to slow down myopic axial eye growth by inducing a relative hyperopia on the peripheral retina (Smith, 2011). These treatments, however, do not prevent or reverse myopia highlighting a gap in knowledge. It is currently not known whether and how optical development of the lens is modulated in coordination with the growing eye. Our data of tight coordination between axial eye length and the lens nucleus localization, which serves as a proxy for optical development, particularly in the regenerating eyes strongly suggests that lens optics are influenced by axial eye length.

Interestingly, the *X. laevis* lens nucleus centralization plateaued at a value of r/a ∼0.08 (Fig. 1A) compared with the 0.0 of the zebrafish (Vorontsova et al., 2018) placing the nucleus slightly more anterior in *X. laevis*. So, while the overall pattern of centralization between the two species was similar, the exact values of lens nucleus location slightly vary. This could be due to species differences in the proportions between the lens diameter to axial eye length diameter, the positioning of the eyes on the animal, the variation of lighting in their environments, or differences in optical requirements. Although *Xenopus* are fully aquatic frogs, it has been shown that adults largely use vision for seeing in air, unlike tadpoles (Chung et al., 1975). At metamorphosis, *Xenopus* eyes become hyperopic in water and emmetropic in air with focal point coinciding with the photoreceptors (Chung et al., 1975). The positioning of eyes at the top of the head after metamorphosis is also suited for vision above their head at the surface or above the water’s surface. These behavioral requirements for vision may require a higher power lens for that can be facilitated by a more anteriorly placed lens nucleus in adults.

Aside from the lens nucleus localization, the gradient of refractive index is the second parameter that contributes to lens power development. Assessment of lens GRIN development and its correlation with centralization of the lens nucleus and axial eye length will determine whether the focal length for focusing light correlates with the retina of the elongating eye.

Our study has shown that the centralization of the lens nucleus is a conserved mechanism of optical development in two aquatic species. This process closely correlates with eye axial length strengthening the idea that these two parameters progress in tight coordination to form a functional optical system. This coordination is key to understanding how normal optics form, and what may go wrong leading to mismatch and refractive error. The use of the lens nucleus as a biomarker of optical development in *Xenopus laevis* and *Danio rerio*, together with the advantages of these species such as molecular modification, live imaging and regeneration can be used as a powerful tool to unravel new mechanisms of normal and pathological optical development and provide new avenues for refractive error prevention.

## Funding

This work was supported by the National Institutes of Health (EY031587, GM146672, HD105600), the Aotearoa New Zealand National Eye Centre fellowship (IV), The Society for Developmental Biology Choose Development! Fellowship (HD105600) (KG) and an NSF REU research fellowship (2244087) (KG).

## CRediT authorship contribution statement

**Karla Garcia:** Writing – original draft preparation, Methodology, Investigation, Visualization, Funding acquisition. **Kelly Tseng:** Writing – review & editing, Supervision, Methodology, Resources, Funding acquisition, Project Administration, Conceptualization. **Irene Vorontsova:** Writing– original draft, Writing – review & editing, Supervision, Methodology, Formal Analysis, Validation, Visualization, Funding acquisition, Conceptualization.

## Acknowledgement

We would like to thank the Tseng lab and the Molecular Vision Research Cluster headed by Prof. Donaldson for helpful comments and suggestions.

## Declaration of competing interest

The authors declare no financial interest.

## Notes

### Competing Interest Statement

The authors have declared no competing interest.

## REFERENCES

Álvarez Hernán, G., de Mera-Rodríguez, J.A., Gañán, Y., Solana-Fajardo, J., Martin-Partido, G., Rodriguez Leon, J., Francisco-Morcillo, J., 2021. Development and postnatal neurogenesis in the retina: A comparison between altricial and precocial bird species. Neural regeneration research 16, 16–20.

Aoki, H., Ogino, H., Tomita, H., Hara, A., Kunisada, T., 2016. Disruption of Rest Leads to the Early Onset of Cataracts with the Aberrant Terminal Differentiation of Lens Fiber Cells. PLOS ONE 11, e0163042.

Beach, D.H., Jacobson, M., 1979. Patterns of cell proliferation in the retina of the clawed frog during development. Journal of Comparative Neurology 183, 603–613.

Cheng, M.H., Tam, C.N., Choy, K.W., Tsang, W.H., Tsang, S.L., Pang, C.P., Song, Y.Q., Sham, M.H., 2016. A γA-Crystallin Mouse Mutant Secc with Small Eye, Cataract and Closed Eyelid. PLOS ONE 11, e0160691.

Chung, S.H., Stirling, R.V., Gaze, R.M., Land, M., Stirling, R.V., 1975. The structural and functional development of the retina in larval Xenopus. Development 33, 915–940.

Danka Mohammed, C.P., Khalil, R., 2020. Postnatal Development of Visual Cortical Function in the Mammalian Brain. Frontiers in Systems Neuroscience Volume 14 -2020.

Faber, J., & Nieuwkoop, P.D. (Eds.), 1994. Normal Table of Xenopus Laevis (Daudin): A Systematical & Chronological Survey of the Development from the Fertilized Egg till the End of Metamorphosis 1st ed. Garland Science.

Fleisch, V.C., Neuhauss, S.C., 2006. Visual behavior in zebrafish. Zebrafish 3, 191–201.

Gaze, R.M., 1959. Regeneration of the optic nerve in Xenopus laevis. Quarterly journal of experimental physiology and cognate medical sciences 44, 290–308.

Gestri, G., Link, B.A., Neuhauss, S.C., 2012. The visual system of zebrafish and its use to model human ocular diseases. Dev Neurobiol 72, 302–327.

Greiling, T.M., Clark, J.I., 2008. The transparent lens and cornea in the mouse and zebra fish eye. Semin Cell Dev Biol 19, 94–99.

Guerin, D.J., Gutierrez, B., Zhang, B., Tseng, K.A.-S., 2025. Notch Is Required for Neural Progenitor Proliferation During Embryonic Eye Regrowth. International Journal of Molecular Sciences 26, 2637.

Hack, S.J., Petereit, J., Tseng, K.A.-S., 2024. Temporal Transcriptomic Profiling of the Developing Xenopus laevis Eye. Cells 13, 1390.

Hamor, R., Whitley, D., 2024. Ocular Physiology and Vision in the Equine Neonate, Equine Neonatal Medicine, pp. 1185-1196.

Ide, C.F., Reynolds, P., Tompkins, R., 1984. Two healing patterns correlate with different adult neural connectivity patterns in regenerating embryonic Xenopus retina. The Journal of experimental zoology 230, 71–80.

Kha, C.X., Guerin, D.J., Tseng, K.A.-S., 2019. Using the Xenopus Developmental Eye Regrowth System to Distinguish the Role of Developmental Versus Regenerative Mechanisms. Frontiers in Physiology Volume 10 -2019.

Kha, C.X., Nava, I., Tseng, K.A., 2023. V-ATPase Regulates Retinal Progenitor Cell Proliferation During Eye Regrowth in Xenopus. Journal of ocular pharmacology and therapeutics : the official journal of the Association for Ocular Pharmacology and Therapeutics 39, 499–508.

Kha, C.X., Son, P.H., Lauper, J., Tseng, K.A., 2018. A model for investigating developmental eye repair in Xenopus laevis. Exp Eye Res 169, 38–47.

Kroger, R.H., 2013. Optical plasticity in fish lenses. Prog Retin Eye Res 34, 78–88.

Land, M.F., Fernald, R.D., 1992. The evolution of eyes. Annual review of neuroscience 15, 1–29.

Leach, L.L., Hanovice, N.J., George, S.M., Gabriel, A.E., Gross, J.M., 2021. The immune response is a critical regulator of zebrafish retinal pigment epithelium regeneration. Proc Natl Acad Sci U S A 118.

Lie, A.L., Pan, X., Donaldson, P.J., White, T.W., Vaghefi, E., 2018. The lens paradox is due to an age-related shift in optical centre of the lens. Investigative Ophthalmology & Visual Science 59, 4698–4698.

Liu, Z., Hamodi, A.S., Pratt, K.G., 2016. Early development and function of the Xenopus tadpole retinotectal circuit. Current Opinion in Neurobiology 41, 17–23.

Mitra, A.T., Womack, M.C., Gower, D.J., Streicher, J.W., Clark, B., Bell, R.C., Schott, R.K., Fujita, M.K., Thomas, K.N., 2022. Ocular lens morphology is influenced by ecology and metamorphosis in frogs and toads. Proc Biol Sci 289, 20220767.

Mutti, D.O., Sinnott, L.T., Lynn Mitchell, G., Jordan, L.A., Friedman, N.E., Frane, S.L., Lin, W.K., 2018. Ocular Component Development during Infancy and Early Childhood. Optometry and vision science : official publication of the American Academy of Optometry 95, 976–985.

Pankhurst, P.M., Eagar, R., 1996. Changes in visual morphology through life history stages of the New Zealand snapper, Pagrus auratus. New Zealand Journal of Marine and Freshwater Research 30, 79–90.

Pérez-Montes, C., Hernández-García, R., Jiménez-Cubides, J.P., DeOliveira-Mello, L., Velasco, A., Arévalo, R., García-Macia, M., Santos-Ledo, A., 2026. Zebrafish optic nerve regeneration involves resident and retinal oligodendrocytes. Neural Regeneration Research 21, 811–820.

Rio-Tsonis, K.D., Tsonis, P.A., 2003. Eye regeneration at the molecular age. Developmental Dynamics 226, 211–224.

Sivak, J.G., 1988. Optics of Amphibious Eyes in Vertebrates. Springer New York, New York, NY, pp. 467–485.

Smith, E.L., 3rd, 2011. Prentice Award Lecture 2010: A case for peripheral optical treatment strategies for myopia. Optometry and vision science : official publication of the American Academy of Optometry 88, 1029–1044.

Suetsugu-Maki, R., Maki, N., Nakamura, K., Sumanas, S., Zhu, J., Del Rio-Tsonis, K., Tsonis, P.A., 2012. Lens regeneration in axolotl: new evidence of developmental plasticity. BMC Biology 10, 103.

Summers, J.A., Schaeffel, F., Marcos, S., Wu, H., Tkatchenko, A.V., 2021. Functional integration of eye tissues and refractive eye development: Mechanisms and pathways. Exp Eye Res 209, 108693.

Tripathi, R.C., Tripathi, B.J., 1983. Lens morphology, aging, and cataract. Journal of gerontology 38, 258–270.

Troilo, D., Smith, E.L., 3rd, Nickla, D.L., Ashby, R., Tkatchenko, A.V., Ostrin, L.A., Gawne, T.J., Pardue, M.T., Summers, J.A., Kee, C.S., Schroedl, F., Wahl, S., Jones, L., 2019. IMI - Report on Experimental Models of Emmetropization and Myopia. Invest Ophthalmol Vis Sci 60, M31–M88.

Tseng, A.-S., 2017. Seeing the future: using Xenopus to understand eye regeneration. genesis 55, e23003.

Tsonis, P., Rio-Tsonis, K., Washabaugh, C., 1993. Analysis of the mutant axolotl short toes. Progress in clinical and biological research 383A, 171–179.

Viet, J., Reboutier, D., Hardy, S., Lachke, S.A., Paillard, L., Gautier-Courteille, C., 2020. Modeling ocular lens disease in Xenopus. Dev Dyn 249, 610–621.

Vorontsova, I., Gehring, I., Hall, J.E., Schilling, T.F., 2018. Aqp0a Regulates Suture Stability in the Zebrafish Lens. Invest Ophthalmol Vis Sci 59, 2869–2879.

Wan, J., Goldman, D., 2016. Retina regeneration in zebrafish. Current opinion in genetics & development 40, 41–47.

Wang, K., Vorontsova, I., Hoshino, M., Uesugi, K., Yagi, N., Hall, J.E., Schilling, T.F., Pierscionek, B.K., 2020. Optical development in the zebrafish eye lens. FASEB J 34, 5552–5562.

Wen, L., Shi, Y.-B., 2016. Regulation of growth rate and developmental timing by Xenopus thyroid hormone receptor α. Development, Growth & Differentiation 58, 106–115.

